# NMDAr Blocking by MK801 Alters Hippocampal and Prefrontal Cortex Oscillations and Impairs Spatial Working Memory in Mice

**DOI:** 10.1101/2021.09.22.461383

**Authors:** P. Abad-Perez, F.J. Molina-Payá, L. Martínez-Otero, V. Borrell, R.L. Redondo, J.R. Brotons-Mas

## Abstract

Abnormal NMDAr function has been linked to rhythmopathies, psychosis, and cognitive dysfunction in schizophrenia (SCZ). Here, we investigate the role of NMDAr hypofunction in pathological oscillations and behavior. We implanted mice with tetrodes in the dorsal hippocampus and medial prefrontal cortex (mPFC), administered the NMDAr antagonist MK801, and recorded oscillations during spontaneous exploration in an open field and in the y-maze spatial working memory test. Our results show that NMDAr blockade increased locomotor activity, impaired spatial working memory, and disrupted the correlation between oscillations and speed of movement, which is crucial for internal representations of distance. In the hippocampus, MK801 increased gamma oscillations and disrupted theta/gamma coupling. In the mPFC, MK801 increased the power of theta and gamma, generated high-frequency oscillations (HFO 155-185 Hz), and disrupted theta/gamma coupling. The performance of mice in the spatial working memory version of the y-maze was strongly correlated with CA1-PFC theta/ low gamma co-modulation. Thus, theta/gamma mediated by NMDAr function might be essential to explaining several of SCZ’s cognitive symptoms and might be crucial to explaining hippocampal-PFC interaction.

**Significance Statement:** NMDAr hypofunction might be the basis of cognitive symptoms and oscillopathies found in SCZ. In this work, we aimed to understand this link further. We found that NMDAr hypofunction altered theta/gamma co-modulation in the hippocampus and the PFC, explaining spatial working memory deficits.

## Introduction

Schizophrenia is a disabling disorder characterized by negative, positive, and cognitive symptoms, e.g., lack of social interest, psychosis, and working memory deficits (Owen et al., 2016). A dominant theory about SCZ’s etiology is the dopaminergic hypothesis, which posits that this disorder is caused by dopamine imbalance. This theory is supported by the therapeutic effects of anti-dopaminergic drugs (Meltzer and Stahl, 1976; Ban, 2007). However, neuroleptic medications fail to treat negative and cognitive symptoms, suggesting that multiple mechanisms are at play in SCZ (Bowie and Harvey, 2006; Kirkpatrick et al., 2006; James et al., 2018). The NMDAr hypofunction hypothesis of SCZ may offer a complementary explanation of the etiology of this disorder, including its cognitive symptoms. NMDAr antagonists can induce SCZ-like symptoms in patients and healthy subjects (Malhotra et al., 1997; Adler et al., 1998). Schizophrenic patients showed a reduced expression of NMDAr in the dorsolateral prefrontal cortex (DLPFC), a brain region strongly involved in working memory and executive function, impaired in SCZ (Beneyto and Meador-Woodruff, 2008; Weickert et al., 2013). NMDAr hypofunction seems to dysregulate dopamine levels (indirectly increasing striatal dopamine levels), linking NMDAr function with dopamine imbalance (Wȩdzony et al., 1993; Vollenweider et al., 2000; Del Arco et al., 2008). Additionally, animal models of SCZ based on NMDAr hypofunction successfully mimic the positive, negative, and cognitive symptoms as well as structural brain changes observed in schizophrenic patients (Olney et al., 1999; Eyjolfsson et al., 2006; Abekawa et al., 2007; Adell et al., 2012; Jodo, 2013; Murueta-Goyena et al., 2017). Finally, the pharmacological blockade of NMDAr produces aberrant oscillatory activity and interneuron dysfunction, also associated with SCZ (Hertzman et al., 1990; Kristiansen et al., 2006; Hakami et al., 2009).

The deletion of NMDAr from parvalbumin-positive interneurons (PV^+^) in rodent models triggers cognitive symptoms recapitulating many of those observed in SCZ (Weickert et al., 2000; Belforte et al., 2010; Korotkova et al., 2010; Mena et al., 2016). Dizolcipine, or MK801, is an NMDAr antagonist that non-competitively blocks NMDAr’s voltage-dependent ion channels (Huettner and Bean, 1988). It is commonly used as a model of SCZ in rodents as it reproduces its cognitive deficits, gamma power alterations, and the emergence of high-frequency oscillations (HFO) across different brain regions. At the cellular level, NMDAr hypofunction induced by MK801 administration reduces the firing of GABAergic interneurons, increasing the firing rate of excitatory pyramidal neurons (Jackson et al., 2004; Homayoun and Moghaddam, 2007). Therefore, the NMDAr hypofunction hypothesis offers a possible mechanism that links the cognitive deficits, the aberrant oscillatory activity, and the interneuron dysfunction observed in SCZ, while at least partially explaining some of the dopamine changes typically observed in patients (Uhlhaas and Singer, 2010; Lewis et al., 2012; Moghaddam and Javitt, 2012).

Despite the accumulation of data regarding the role of NMDAr hypofunction at the level of cellular and synaptic mechanisms, its relationship to cognition remains poorly understood. Here, we aimed to understand how NMDAr blockade by MK801 can affect oscillatory activity, behavior, cognition, and the dialog between the dorsal and intermediate CA1 hippocampal region and the prefrontal cortex (PFC), describing the oscillatory changes associated with behavioral alterations. These hippocampal regions are connected directly and indirectly with the PFC (Hoover & Vertes, 2007), and their interaction is crucial for executive function and SCZ (Jones and Wilson, 2005; Sigurdsson and Duvarci, 2016). We were especially interested in determining the oscillatory correlates involved in spatial cognition and decision-making in spatial working memory. For this reason, we recorded simultaneously from the intermediate CA1 and the PFC to study the effect of MK801-mediated NMDAr blockade on oscillatory activity, behavior, and spatial working memory performance. We expected to find changes in the gamma band and distortion of the hippocampus/PFC interaction associated with working memory deficits. Our results were consistent with these predictions. In addition, NMDAr blockade generated area-specific oscillatory changes in the hippocampus and the PFC, and altered the relationship between oscillatory activity and motor behavior.

## Methods

### Subjects

Male wild-type mice C57, N=9, age p60 to p90 supplied by the UMH “Servicio de Experimentación Animal (SEA)” were used. Mice were maintained on a 12 h light/dark cycle with food and water available *ad libitum* and were individually housed after electrode implantation. Experimental procedures were approved by the UMH-CSIC ethics committee and the regional government and met local and European guidelines for animal experimentation (86/609/EEC).

### In vivo recordings on freely moving mice

Microdrives, (Axona ltd) mounting four tetrodes (12 μm tungsten wire, California Fine Wire Company, Grover Beach, CA, USA) were implanted under isoflurane anesthesia (1.5%) while buprenorphine (0.05mg/kg, s.c.) was supplied as analgesia. Craniotomies were performed above the hippocampus and medial prefrontal cortex (mPFC), targeting the hippocampal CA1 region (AP: −2-2.5, M-L: 1.2 V: 0.6 mm) as well as the prelimbic region of the mPFC (AP: 1-2, M-L: 0.5-1, V: 1.2-1.9 mm). Animals were left to recover for at least seven days after surgery (Brotons-Mas et al., 2010; Brotons-Mas et al., 2017; del Pino et al., 2013), see Fig 1A.

**Figure 1.**
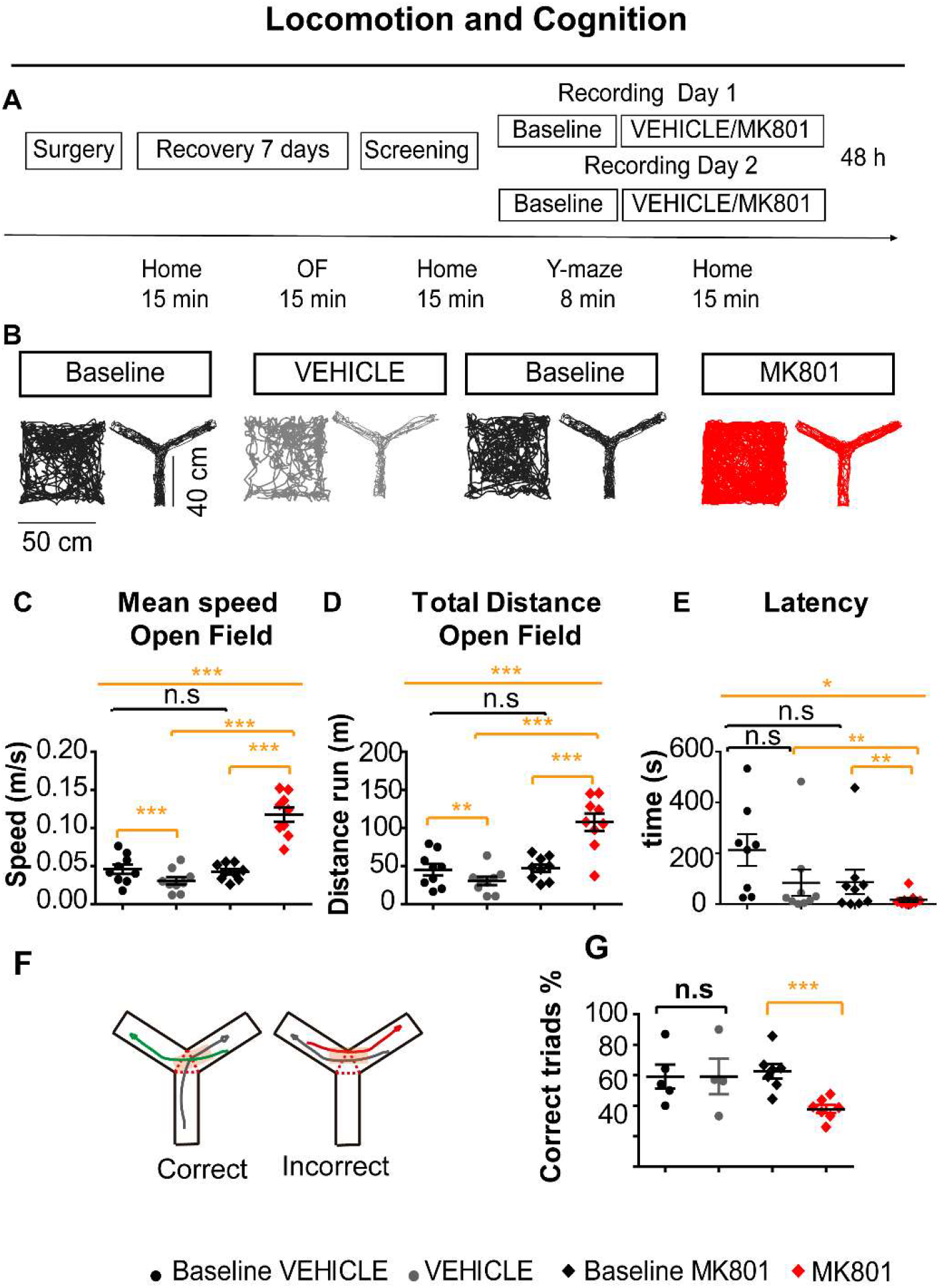
MK801 increased exploration and locomotor behavior. A. Experimental protocol, open field (OF). B. Representative tracking for the baseline, vehicle, and MK801 condition in the open field and y-maze experiment. C. Mean speed was reduced during the vehicle condition, possibly as result of habituation to the environment, but was increased after MK801 administration. D. Total distance run was augmented during the MK801 condition. E. Latency to visit the center of the open field, only 8 points are displayed as one of the animals never visited the center. Note that the behavior across days in the baseline conditions was very similar indicating behavioral consistency across days. F Examples of correct and incorrect trials. G. Lower performance in the y-maze spatial working memory task after NMDAr blockade.

### Data acquisition

Electrophysiological recordings were obtained using a 16-channel headstage, gain x1 (Axona Ltd, UK), (Brotons-Mas et al., 2010; Brotons-Mas et al., 2017; del Pino et al., 2013). Signals were amplified (400 to 1000x), bandpass filtered (0.3 Hz to 24 kHz, DacqUSB system, Axona, UK) and recorded at 48 kHz/24-bit precision. For Local field potential recordings (LFP), tetrodes were lowered to reach specific stereotaxic coordinates in the mPFC. Unit activity, ripples (high-frequency events 150-200Hz), and theta power were used as electrophysiological landmarks to determine electrode location in the CA1 pyramidal layer. Once tetrodes were in the target areas, we started the recording protocols. This part of the procedure was implemented for several days to ensure recording stability.

The mouse trajectory was recorded by a video-tracking system (Axona) that detected the position of an infrared light-emitting diode on the headstage. Animal head position information was stored at 50 Hz in X, Y coordinates, with a spatial resolution of 322 pixels / m and timestamped relative to the electrophysiological recordings. Position data was used to calculate the speed of movement and all the exploratory related variables. For this, custom-made and adapted Matlab scripts were used, see Fig 1B.

### Power spectrum analysis

Local field potentials were analyzed using custom-written Matlab codes (Mathworks, Natick, MA, USA). Raw recordings were FIR filtered (<1.2 kHz) and downsampled to 2.4 kHz, as in previous work (del Pino et al., 2017). Data obtained in open field recordings was used to characterize oscillatory activity. Running speed was computed based on the animal’s position. Spectral power was calculated (in decibels 10log10) using the Thomson multi-taper method included in Chronux signal processing toolbox (Bokil et al., 2010). The spectrogram was plotted to visualize the power spectrum in the time domain notch filter (50 Hz and 100 Hz) was implemented to eliminate electrical noise. We excluded from analysis high-frequency noise (135-145 Hz) associated with chewing artifacts, and we eliminated it from the periodogram plots. The functional connectivity between the hippocampus and the prefrontal cortex was measured by calculating frequency coherence. This measurement describes the degree of co-occurrence of different frequencies in the activity recorded across different areas.

To establish the correlation between oscillatory activity and the speed of movement, we segmented the signal in non-overlapping windows of one second. Then, we calculated the power spectrum and obtained the theta (4-12Hz) / delta (<3.9Hz) ratio to eliminate epochs contaminated by mechanical artifacts, such as head bumping against the wall etc. We calculated the power spectrum for theta type I (6.5-12Hz), linked to motor behavior. In addition, we obtained the power values for low (30-65 Hz) and high gamma (66-130 Hz) bands. Complementary, we calculated oscillatory power for epochs in which locomotor activity was <5 cm/s. The power of theta and co-modulated gamma are known to be higher during epochs of higher speeds. Therefore, the consideration of velocities can help to control for this confounding variable.

Then, to control for volume propagation in the PFC, we performed a bipolar derivation, subtracting the signal obtained from two channels in different tetrodes implanted in the PFC and we performed the same LFP analysis previously described.

### Gamma modulation index (GMI)

The gamma modulation index was calculated as previously described (Korotkova et al., 2010). In brief, the raw LFP was filtered in the theta band (4-12 Hz) and in different higher frequency intervals: low gamma (30-65 Hz), high gamma (66-130), and high-frequency oscillations (155-185 Hz). The gamma and HFO envelopes were calculated to determine the power of the signal across the different theta cycles; the maximum and minimum values were used to calculate the gamma index of modulation (GMI).

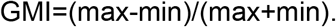

Also, hippocampal theta/ PFC gamma bands co-modulation was calculated to measure region interaction using the same procedure.

### Behavioral Protocol and drug administration

Once animals recovered from surgeries, tetrodes aiming to the CA1 and the PFC were lowered during several days until target coordinates were reached. Experiments begun with a baseline condition. This served to calculate ratios of LFP power to determine change due to drug administration controlling for possible electrode position drifting across days.

Each baseline and drug condition consisted of several phases. First, recordings were performed in the animal’s home cages for several minutes to ensure that electrodes were placed in the CA1 pyramidal layer. We recorded LFPs during spontaneous exploration in an open field of 50 x 50 cm. Subsequently mice were left to rest for five minutes in their home cage. After this period, mice underwent testing in the y-maze for 8 minutes and left to rest in their home cage. Once the baseline was finished, animals were left to rest for 20 min in their home cage. At that point, mice were injected subcutaneously with MK801 (0.075 mg/kg) diluted in 0.3% tween80 in saline or with the vehicle solution, consisting of diluted in 0.3% tween80 in saline. Behavioral testing and recordings followed the same sequences previously explained. Recordings were initiated 20 min after injection and experiments performed within the temporal window of MK801 bioavailability (Wegener et al, 2011).

Drugs were administered following a counterbalanced scheme across alternate days. This way each mouse either received MK801 or vehicle injections on day one, and the complementary treatment on day two. Experiments were spaced at least 48h, see Fig 1A.

### Working memory in the y-maze

We used the y-maze to measure spatial working memory. This test is a reliable approach for cognitive testing and provides a measurement of spatial working memory based on the ethology of mice (Miedel et al., 2017). Testing was carried out on a transparent Y-shaped maze (40 x 10 cm per arm). Mice were placed at the end of the central arm and allowed to explore the maze for 8 minutes. The sequence of arm entries was obtained from the x,y position coordinates and automatically analyzed using custom-made scripts in Matlab. Correct alternations were those in which the animal went into an arm not visited recently, e.g., correct visit sequence: arm 1-2-3. We considered incorrect alternations when animals repeated the same arm in a sequence of trials, e.g. arm 1-2-1. The final score was calculated as the percentage of correct triads using the formula below:

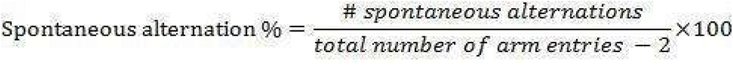

We searched for different electrophysiological biomarkers of spatial working memory to investigate normal and pathological mechanisms during memory performance in the y-maze. To this end, we analyzed LFP epochs of 1s duration, half a second before the mice entered the decision area of the y-maze (the central triangle between the three arms) and half a second later. We only included experiments consisting of at least nine entries, ensuring the sampling of a minimum of LFP epochs. Finally, we included five animals in the vehicle condition and seven in the MK801 experiments, excluding four and two respectively. As animals explored much less in the vehicle condition, we centered our analysis on the baseline vs. MK801 conditions, focusing on the LFP obtained in these sets of experiments and including a final N=7 mice.

### Anatomical verification of electrode location

After completing experiments, animals were deeply anesthetized using sodium pentobarbital (Euthanasol, 1 mL/ kg, i.p.) and transcardially perfused with saline and 4% paraformaldehyde; the brains were removed, sliced in 40 μm sections, and prepared for Immunohistochemistry against DAPI, NeuN (Merck,ltd) and GFAP (Adigene technologies) for electrode localization.

### Statistical analysis

Variables distribution was tested for normality using Kolmogorov-Smirnov test. To determine statistically significant differences across conditions, parametric and nonparametric tests were used. ANOVA repeated measures was used for normally distributed variables and Greenhouse-Geisser correction applied when sphericity criteria was not met. This correction allows for a more conservative interpretation of the results after adjusting the degrees of freedom. Partial eta squared (ɲ^2^) was calculated as an estimation of the size effect and the statistical power reported. The Friedman test was used for non-normally distributed variables. Then, paired comparisons between specific conditions were tested using t-test for repeated measures and non-parametric Wilcoxon test when appropriate. In the case of correlations, normality was verified and the parametric Pearson coefficient were calculated. All descriptive values were expressed as mean ± the standard error (S.E.). Statistical significance was set at p< 0.05, and Bonferroni corrections were used when necessary. All statistical analysis was performed using SPSS (IBM, V27) software package.

## Results

### MK801 generated hyperlocomotion and impaired spatial working memory in the y-maze

SCZ and other psychotic disorders are characterized by positive and cognitive symptoms, e.g. hyperlocomotion and working memory deficits. For this reason, we investigated if the blockade of NMDAr by MK801 could induce these symptoms in normal, wild-type, mice. First, we observed that mean speed, total distance, and latency to visit the center of an open field during exploratory behavior increased after MK801 administration (*speed*, f_(1,11)_= 66.296, p<0.0001, ɲ^2^=0.892, β=1; *total distance*, f_(3,24)_=43.371, p<0.0001; ɲ^2^=0.844, β=1; β=1; *latency*, *χ*^2^_(3)_=10.950, p=0.012, *Friedman test*). Specific analysis (Fig 1B to 1E) demonstrated that MK801 increased motor activity above baseline condition and vehicle condition (baseline vs MK801; *speed*, t_(8)_= −8.6, p=0.0001; *distance*, t_(8)_= −6.918, p=0.0009; Bonferroni corrections p<0.012 *Wilcoxon test; speed*, t_(8)_= −8.9, p=0.0001; *distance*, t_(8)_= −7.19, p=0.001). Interestingly, vehicle administration marginally decreased exploration behavior (t_(8)_= 4.39, p=0.0019; *distance*; t_(8)_= 3.39, p=0.008), probably an effect of familiarity not observed after MK801.

Then we investigated spatial working memory in the Y-maze, (Fig 1F and 1G). NMDAr blockade increased locomotion and reduced performance in the y-maze (*% of alternation*, 62.58±4.8 vs. 37.38±2.56; t_(6)_= 4, p=0.007), with no effects of vehicle (59.08±7.9 vs. 57.34±9.2; t_(4)_= 0.125, p=0.907).

Therefore, NMDAr blockaded by MK801 administration reliably increased locomotor activity and impaired the performance of mice in the y-maze spatial working memory test, replicating positive and cognitive symptoms observed in SCZ.

### Effect of MK801 in oscillatory activity during spontaneous exploration

Theta and gamma oscillation in HPC and PFC are strongly linked with the cognitive demands of spatial working memory tasks (Sigurdsson et al, 2010) and is critical for the generation of spatial memory (Richard et al., 2013; Long et al., 2014; Young et al., 2020; Long et al., 2014; Kramis et al., 1975). Alteration of these dynamics is a landmark in SCZ (Uhlhaas and Singer, 2010). For this reason, we investigated if NMDAr blockade by MK801 alters brain oscillations.

First, we explored the relationship between the speed of movement and theta (6.5-12Hz), and low and high gamma activity (30-65 Hz and 66-130Hz)) in the hippocampal region CA, (Fig 2A to 2C). We found a significant reduction in the correlation between high gamma oscillations and the speed of movement (theta vs speed: f_(1,15)_= 3.087, p=0.59, ɲ^2^=0.382, β=0.6; High gamma: *baseline vs. vehicle*, *r*=,0.52±0.03 vs. 0.59±0.011; t_(8)_= −3.179, p=0.013; *baseline vs. MK801, (r)*, 0.55±0.03 vs. 0.37±0.05; z=-2.547, p=0.01, *Wilcoxon; vehicle vs. MK801*, t_(8)_= 5.598, p=0.0005).

**Figure 2.**
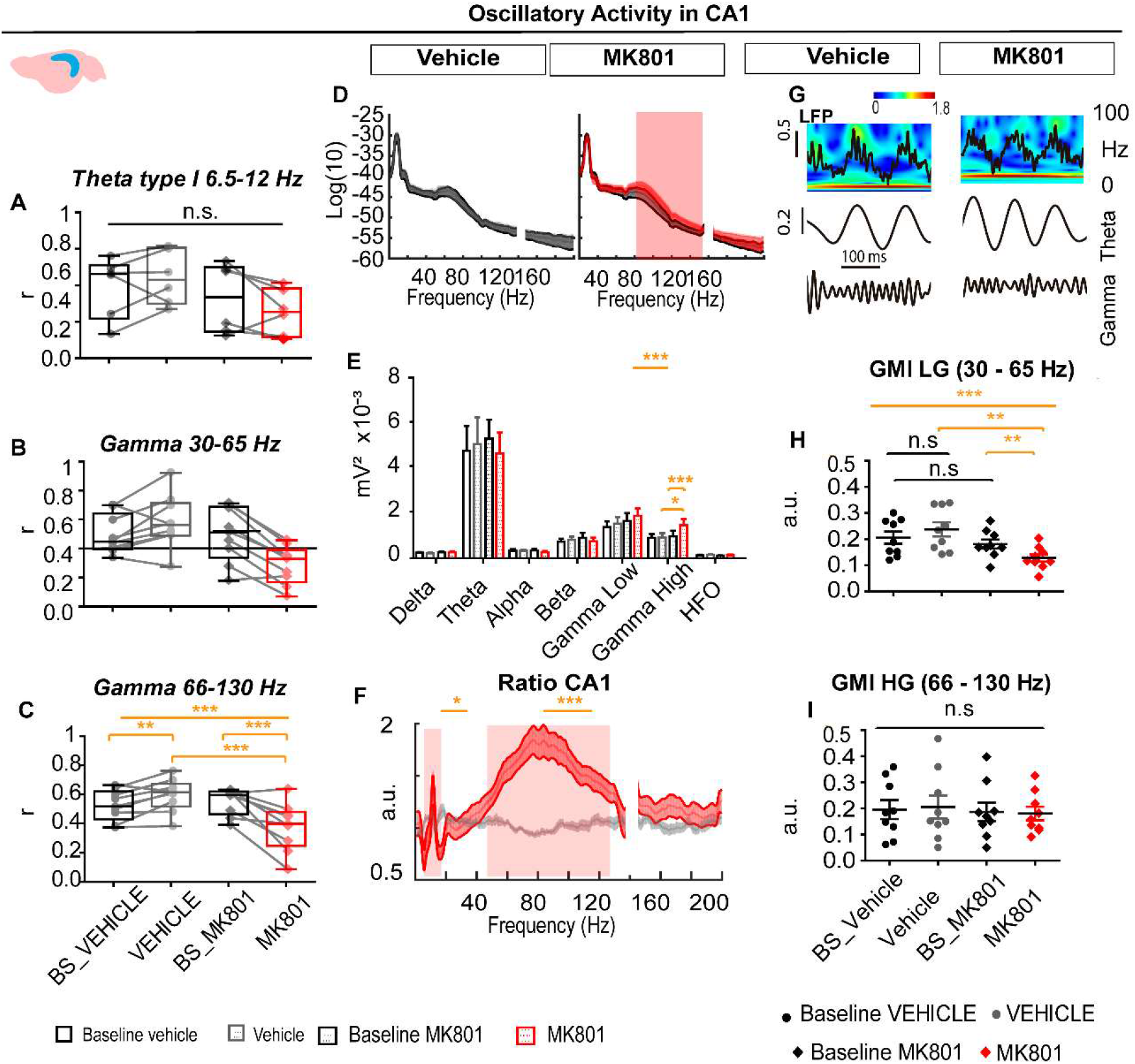
MK801 administration disrupted normal hippocampal oscillatory activity. A. Mean correlation values between the speed of movement, theta type I (6.5-12HZ), low gamma and high gamma. Analysis of theta correlations only included animals showing significant correlations, (n=6). Results indicated that theta /speed correlation decreased not reaching significant values after MK801 administration. B. Low gamma presented a poor correlation with the speed of movement and no further statistical test was performed. C. High gamma presented a robust correlation with the speed of movement (n=9). High gamma showed a significant increase in the vehicle condition and a reduction during the MK801 experiment. D and E. Power spectrum and mean values for each band obtained in the baselines, vehicle, and MK801 conditions. Significant differences were found for the high gamma band in the MK801 vs. its baseline and the vehicle condition. E. Ratio of power change obtained between baseline conditions vs. vehicle and MK801. Significant differences were mostly consistent with mean power value analysis. Shadowed regions in D and F indicate bands in which significant differences were found. Note how the ratio of change between conditions followed a flat line across the frequency spectrum for the vehicle experiment, while in the MK801 the ratio presented specific peaks at different bands. G Representative local field potential, wavelet, and filtered signal for theta and gamma in CA1 during the baseline and after MK801 administration. Wavelet heat map scales expressed in standard deviations revealed the dominant band at each theta cycle. H. Theta/gamma co-modulación in the CA1 region showed significant differences due to MK801 administration. I. CA1 Theta/high gamma was not significantly modified by MK801. Gamma modulation index (GMI), low gamma (LG), high gamma (HG).

Thus, NMDAr blockade by MK801 altered the relationship between CA1 oscillatory activity (66-130Hz) and the speed of movement. There are multiple reports of gamma activity being specifically altered in SCZ and after NMDAr blockade (Hertzman et al., 1990; Kristiansen et al., 2006; Hakami et al., 2009).To analyze these effects further, we considered only epochs of activity in which the mice were moving above 5 cm/s during at least 2 seconds, thus involving linear displacements. This strategy allows us to compare oscillatory activity during similar behavioral episodes and therefore to control for the MK801 effects in locomotor activity (Fig 2D, 2E). We observed a significant augmentation of the high gamma band after MK801 administration (*baseline vs MK801*, z=-2.666, p=0.008; *vehicle vs MK801*, z=-2.310, p=0.021, *Wilcoxon; Bonferroni correction for two comparisons p<0.025*). Then we calculated the ratio of power change for all frequencies baseline/drug (Fig 2F). MK801 effects were more robust for the gamma band (*30-130 Hz*, 1.07±0.06 vs. 1.4±0.08, z=-2.31 p=0.021), but we also found differences in slower frequency bands, including beta, delta and theta (au, *3-9Hz*, 1.04±0.03 vs. 0.79±0.0, t_(8)_=3.058, p=0.016; *10-12Hz*, 1.08±0.06 vs. 1.3 ±0.13 t_(8)_=-2.43, p=0.041; 13-20Hz, 1.15±0.07 vs. 0.77±0.03 z=-2.666 p=0.008). These results indicated that a set of frequencies, including gamma, theta, beta and alpha, changed when the baseline vehicle/vehicle ratio was compared vs. the baseline MK801/MK801 ratio. Interestingly, of all these bands, only theta (10-12 Hz) and gamma band (30-130 Hz) were augmented in their power ratio, while the rest were diminished by MK801.

As expected, from the loss of correlation between speed and gamma power (see Fig. 2B), low and high gamma oscillations were also increased at low speeds (below 5 cm/s, including slow displacements or head movements; z=2.547, p=0.011; z=2.666, p=0.008, *Wilcoxon*) after MK801. Therefore, MK801 general effects on gamma power was independent of the speed of movement.

MK801 specifically increased power in theta and gamma frequency bands, which temporal dynamics have been extensively associated with spatial working memory and sensory processing (Buzsaki et al., 2003; Tamura et al, 2017; Valero and de la Prida, 2018). Based in previous work (Korotkova et al., 2010), we calculated a modulation index (GMI) for theta/low gamma at 30-65 Hz, theta/high gamma, 66-130 Hz (Fig 2G). We found a significant reduction of the theta/low-gamma GMI after MK801 administration (baseline MK801 vs. MK801; t_(8)_=3.982, p=0.004; *vehicle vs. MK801* t_(8)_=4.033, p=0.004; Bonferroni corrections, p=0.0125).

Our results indicated that oscillations in the hippocampal region were altered in different ways. First, the relationship between locomotion and oscillatory activity was impaired. Secondly, the power of the gamma band was augmented. Third, the coordination of the theta and gamma oscillations was diminished. All these elements are critical for cognitive processing involving the activity timing of multiple groups of neurons and might be relevant for cognitive deficits observed in SCZ.

Next, we focused in our simultaneously recorded tetrodes from PFC, a main output region of the hippocampal formation (Hoover & Vertes, 2007), and whose function seems to be critically compromised in SCZ. We used for the PFC recordings the same rationale than in the CA1 region. Repeated measures analysis revealed significant changes in different oscillatory bands: theta, alpha, high gamma and HFO, (see Fig 3A). Post-hoc comparison demonstrated that theta, alpha and high gamma activity was significantly higher in the MK801 condition (theta: *baseline vehicle vs. vehicle;* t_(8)_=-0.547 p=0.599; *baseline MK801 vs. MK801*, t_(8)_=-3.931 p=0.004; *vehicle vs. MK801;* t_(8)_=-3.837 p=0.005.; *Alpha: baseline MK801 vs. MK801*, t_(8)_=-3.197 p=0.013; *vehicle vs. MK801*, t_(8)_=-2.420 p=0.042; *Bonferroni correction p<0.016;* (baseline *MK801 vs. MK801*,z=-2.192, p=0.028). We also observed that HFOs increased after MK801 administration (z=-2.310, p=0.021). As previously described, we performed the ratio comparisons, (Fig 3B). Results indicated that frequency bands between 0.9 to 16 Hz, including delta, theta and alpha, and at higher extent, HFO were significantly increased in the MK801 condition (t_(8)_=-4.188, p=0.003; *HFO*; t_(8)_=-3.212, p=0.026).

**Figure 3.**
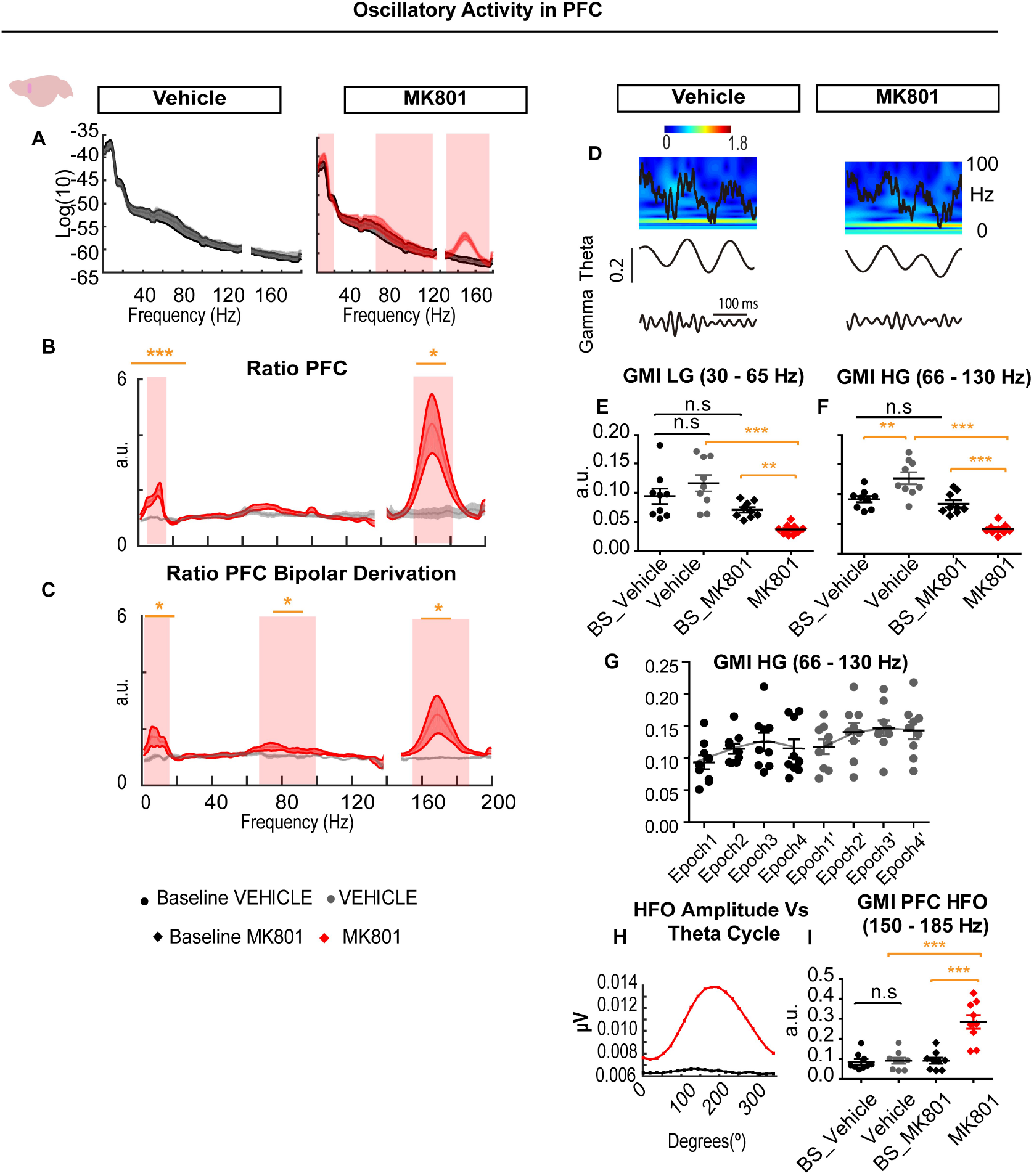
MK801 administration disrupted oscillatory activity in the PFC. A. Power spectrum in the baselines, vehicle and MK801 conditions for monopolar configuration. Note the increase in theta and HFO. B and C. Ratio of power obtained between baseline conditions vs. vehicle and MK801, for monopolar and bipolar derivation analysis. Significant differences were coherent with mean power value analysis for each oscillatory band. Shadowed regions indicate bands in which significant differences were found. D Representative local field potential, wavelet analysis revealing the dominant frequencies, and filtered signal for theta and gamma in the PFC. E. PFC Theta/low gamma was significantly reduced after MK801 administration. F. We observed significant differences between the baseline vs the vehicle condition and a reduction of the GMI after MK801 administration. G. GMI values increased during different temporal epochs, indicating that experience with the spatial context could enhance theta/gamma co-modulation. This could explain the difference observed between the baseline vs. the vehicle condition H. HFO amplitude was modulated by the theta phase. I. Theta/HFO modulation significantly increased after NMDAr blockade.

As part of the recorded LFP in the PFC might be volume propagated, we performed the same ratio analysis after referencing the signal to electrodes located in different tetrodes implanted in the PFC (Fig 3C). We found a very similar pattern to that observed in the single electrode configuration, with same data trend in in the low frequencies, 0.9-16 Hz, but also finding significance differences in the medium-high gamma, 65-100 Hz and in the HFO band (*au, 0.9-16Hz*, 0.99±0.05 vs. 1.5±0.1; t_(8)_=-2.737, p=0.026; *65-100 Hz* 1.05±0.04 vs. 1.28±0.06; t_(8)_=-3.212, p=0.012; *HFO*, 0.9±0.02 vs. 1.8±0.3, t_(8)_=-2.612, p=0.031).

theta/ low gamma and theta/high gamma co-modulation were significantly decreased after MK801 administration and when compared with the vehicle condition (Fig. 3E - F; low gamma: t_(8)_= 9.9599, p=0.001; t_(8)_= 6.015, p=0.001; high gamma: z=2.666, p=0.008). We also observed a significant increase of high/gamma in the vehicle condition (*baseline vs vehicle:* t_(8)_= −3.837, p=0.005; *baseline vs MK801* z=2.666, p=0.008), probably due to habituation, or experience with the spatial context (Tort et al, 2009; Fernandez-Ruiz et al, 2017). For this reason, we obtained the GMI for different epochs of the baseline and the vehicle condition. The observed progressive increase in the GMI during the exploratory task support that the injection of the vehicle did not generate an effect in this variable (Fig. 3G).

The emergence of HFO in the PFC LFP was strongly linked to theta oscillations (Fig 3H - I). We calculated the GMI for the HFO, and observed that the emergence of HFO was linked to an abnormal and significant increase in theta/ HFO modulation (*baseline vs. vehicle,;* z= −1.955 p=0.051; *baseline vs. MK801;* t_(8)_= −5.969, p=0.001; vehicle vs. MK801; t_(8)_= −4.603, p=0.002; Bonferroni corrections p<0.0125).

Thus, NMDAr blocking by MK801 affected oscillatory bands in the hippocampus and the prefrontal cortex, increasing gamma oscillatory activity in the hippocampal region and in the PFC, where also, theta, alpha and HFO, were augmented. NMDAr blockade impaired theta-gamma co-modulation in both brain regions and linked NMDAr hypofunction with an oscillatory phenotype associated with SCZ.

A critical aspect of working memory and SCZ is the dialog between the hippocampus and the PFC (Sigurdsson et al, 2010). To investigate this interaction, (Fig 4A-B), we performed paired comparisons for specific frequency bands and found an small albeit significant increase in the delta band in the monopolar and bipolar configuration (monopolar, *delta, r*, 0.60±0.005 vs. 0.63±0.01; t_(8)_= −2.54, p=0.0340; *bipolar*, 0.55±0.006 vs. 0.57±0.005, t_(8)_= −3.01, p=0.009). Similarly, we observed a significant coherence augmentation for the alpha band in the bipolar electrode configuration (*alpha*, *r*, 0.55±0.006 vs. 0.57±0.005, t_(8)_= −3.01, p=0.016) see Fig 4B’. Then we investigated long-range co-modulation of theta/gamma activity, known to be relevant for spatial working memory (Tamura et al., 2017). GMI index between both hippocampal theta/PFC low gamma and high gamma were reduced in MK801 condition (Fig. 4C; *low gamma baseline vs. vehicle;* z=-1.836, p=0.066; *baseline vs. the MK801* low gamma, t_(8)_= 8.763, p=0.001; *vehicle vs MK801* t_(8)_= 5.558, p=0.000) in consonance with previos results (see Fig. 3??). Hippocampal theta/high gamma PFC modulation was marginally increased during the vehicle condition, as observed in PFC, probably due to the repeated experience with the experimental context. On the other hand, we observed no significant differences in the hippocampal theta/PFC HFO modulation (*baseline vs. vehicle HFO*; Z=-1.224, p=0.214; baseline vs. the MK801 HFO; t_(8)_= −2.240 p=0.055).

**Figure 4.**
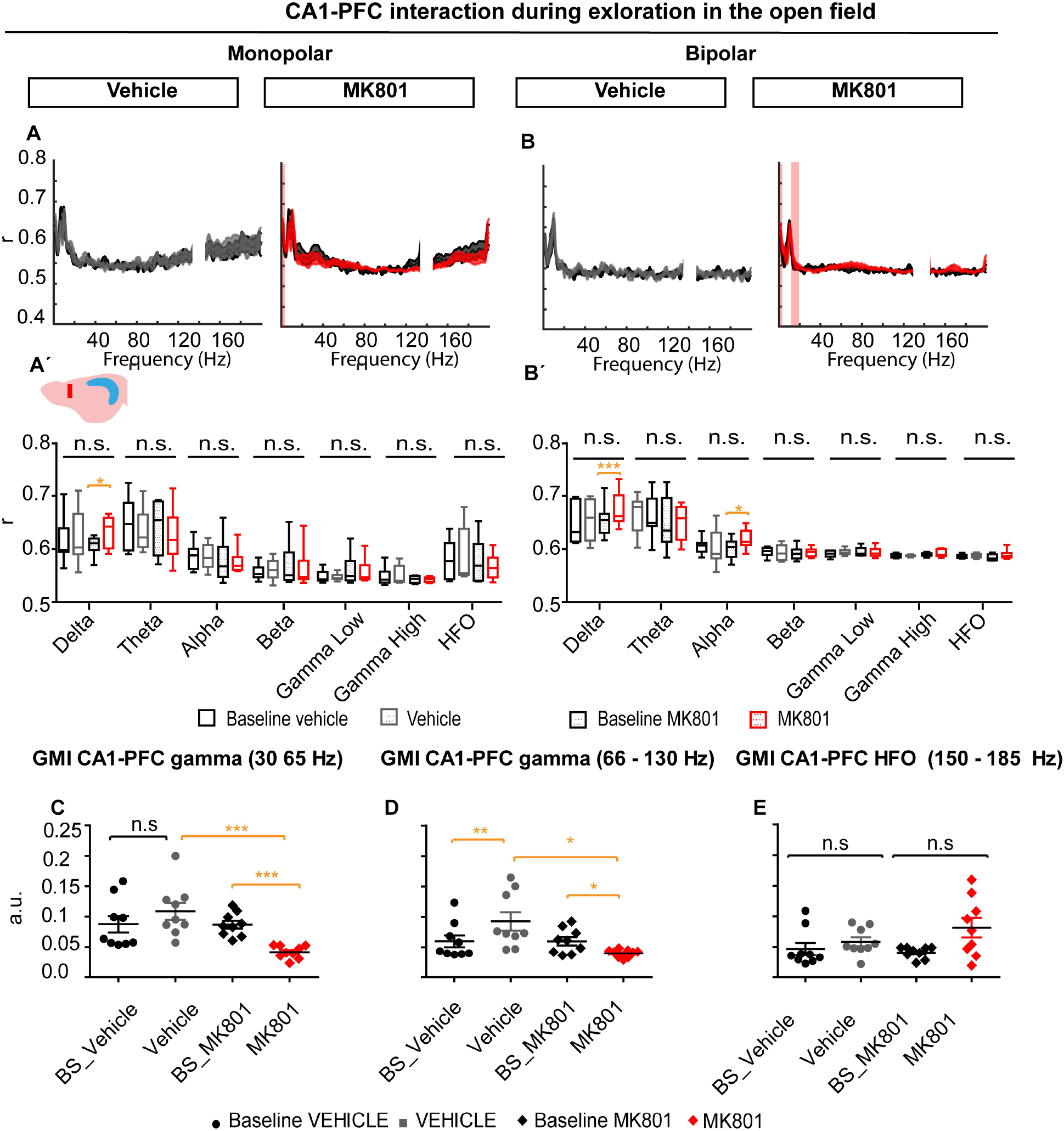
MK801 administration altered CA1-PFC interaction changing theta/gamma co-modulation. A. Coherence values for the baseline vs. vehicle and baseline vs. MK081 obtained in single electrode configuration. A’. Mean coherence values by oscillatory band. B. Coherence values for the bipolar configuration. Shadowed regions indicate significant differences. B’ Mean coherence distribution for different bands. C. Long-range theta/low gamma co-modulation. Significant differences are described by each comparison of interest. D. Theta/high gamma comodulation. E. Theta/HFO co-modulation.

**Figure 5.**
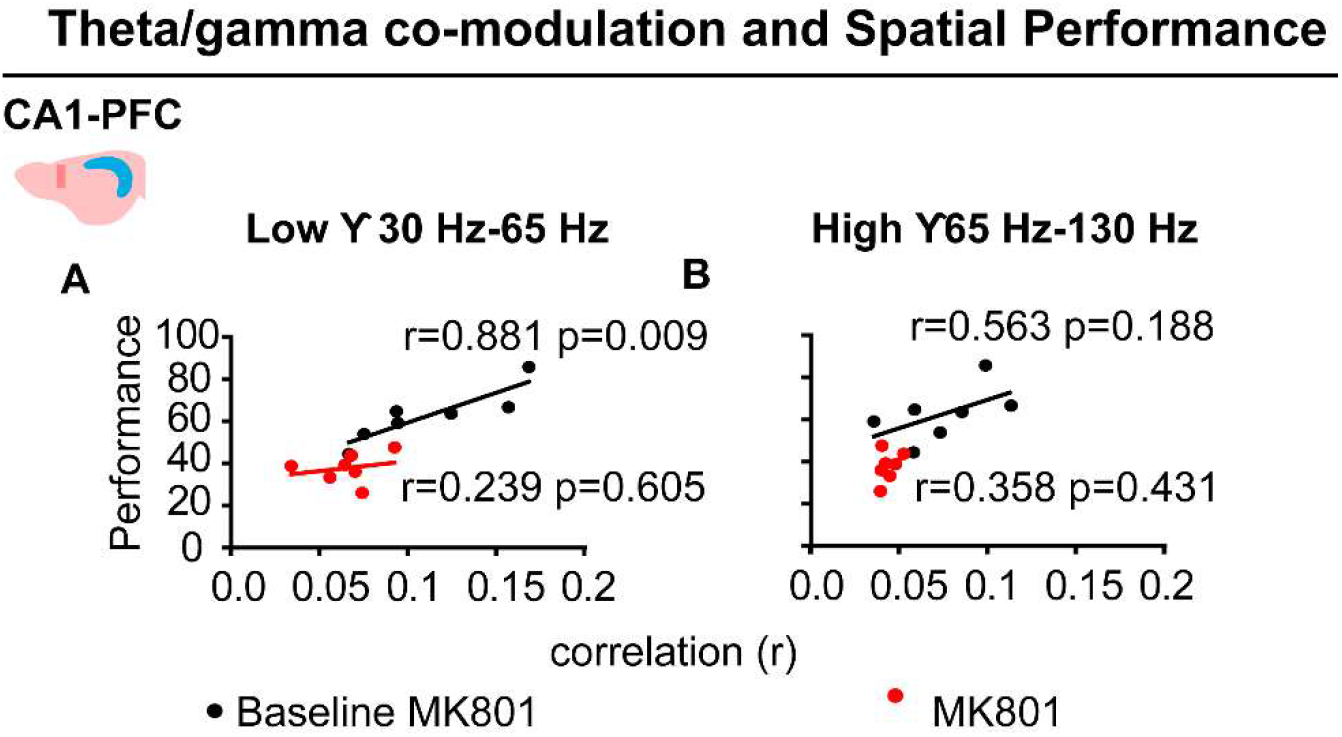
Theta/ gamma co-modulation in CA1 and the PFC correlated with the performance of mice. A. CA1-PFC theta low gamma modulation significantly correlated with the performance of mice in the y-maze. MK801 administration disrupted this correlation. B. Theta/high gamma was significantly correlated with the performance of mice in the Y-maze after MK801 administration.

Therefore, the interaction of the hippocampus and the PFC was altered in specific bands as indicated by the coherence analysis and through the distortion of the theta/gamma co-modulation.

### MK801 disrupted oscillatory activity and cognitive processing in the y-maze

As previously mentioned, there is a relationship between oscillatory activity, cognitive processes and the distortion of these in SCZ (Bygrave et al., 2016, Uhlhaas and Singer, 2010; Sigurdsson et al, 2010). Our results demonstrated that NMDAr administration produced an impairment in working memory in the y-maze. On the other hand, we observed an alteration of oscillatory activity in the hippocampus and the PFC. Thus, we aimed to investigate if there were specific links between the altered oscillatory pattern and performance in the Y-maze, similar to those observed in patients..

We looked at the different oscillatory bands linked to spatial navigation and working memory, especially theta and gamma activity, in the CA1 and PFC, and the interaction between both structures. We observed non-significant differences that could be accounted for by trial type in the oscillatory activity in CA1 or PFC for theta, low, and high gamma oscillations, data not shown. Then, we investigated if the theta / gamma modulation was linked to the performance of mice (Tamura et al, 2017). We could verified that CA1PFC theta/low gamma modulation strongly correlated with the performance Y-maze performance. This correlation was disrupted after MK801 administration. On the other hand, theta/high gamma modulation significantly correlated with the performance of mice after MK801 administration. These results might indicate a specific relationship of low and high gamma oscillations with cognitive process during physiological and pathological conditions.

## Discussion

### NMDAr blockade induced positive-like and cognitive symptoms

Positive symptoms such as hallucinations, delusions, or hyperactivity are one of the hallmarks of SCZ and psychotic episodes (Owen et al., 2016). Throughout our experiments, MK801 induced hyperlocomotion, increasing the speed of movement, exploratory time, and the total number of entries in the y-maze. The augmentation of motor behavior was consistent in the open field and the y-maze and could be interpreted as reproducing SCZ’s positive symptoms. As previously described, this hyperactivity correlates with increased levels of dopamine release in the striatum, possibly mediated by MK801’s action on NMDAr in PV+ interneurons (Belforte et al., 2010; Carlén et al., 2012; Bygrave et al., 2016). On the other hand, animal behavior was very consistent across baseline conditions across different days. Another of the hallmarks of SCZ is its associated cognitive deficits, especially those related to executive function, which are accompanied by aberrant oscillatory activity (Uhlhaas and Singer, 2010; Lewis et al., 2012). In agreement with previous work (Bygrave et al., 2016), NMDAr blockade impaired spatial working memory, a form of executive function, in the y-maze memory test (Callicott et al., 2000; Miedel et al., 2017). This memory deficit was associated with a significant decrease in the long-range theta/gamma co-modulation, indicating a distorted dialog between the hippocampus and the PFC.

Therefore, NMDAr hypofunction might be central for the different positive and cognitive symptoms observed in SCZ either directly or indirectly, e.g., mediating the activity of dopamine levels and altering the dialog between different brain regions.

### NMDAr blockade altered the relationship between hippocampal oscillations and locomotion

In physiological conditions, different aspects of theta and gamma oscillations correlate with the speed and the acceleration of movement, mediating spatial cognition (Whishaw and Vanderwolf, 1973; Chen et al., 2011; Long et al., 2014; Kropff et al., 2021). This is further supported by the fact that the association strength between theta and speed predicts the performance in spatial tasks (Richard et al., 2013.; Young et al., 2020). We observed that MK801 disturbed the normal association between speed of movement and oscillations, especially with respect to high gamma activity and marginally theta oscillations.

By disrupting the relationship between oscillatory activity and motor behavior, NMDAr hypofunction might cause the loss of spatial information in place cells, the impairment of the internal odometer, and therefore the integration of egocentric and allocentric information. Also, oscillatory changes could generate functional disconnections between different brain regions (Jackson et al., 2004; Terrazas et al., 2005; Inostroza et al., 2013; AA, Housh et al, 2014).

### NMDAr blockade differentially altered the oscillatory activity in the hippocampus and PFC

Previous work indicated that NMDAr blockade produced similar effects across different brain regions. However, in our experiments, MK801 produced different oscillatory profiles in the hippocampus and PFC. While oscillatory changes in the CA1 were characterized by changes in the gamma power, oscillatory activity under MK801 in the PFC was characterized by an increase in theta, beta, high gamma, and HFO. Unlike in previous reports (Olszewski et al., 2013), changes in the PFC oscillations, including theta, gamma, HFO, etc., were maintained after bipolar derivation, as revealed by the analysis of ratios, (baselines/experimental condition). Reductions of hippocampal theta have been described in previous work (Hargreaves et al., 1997; Kittelberger et al., 2012; Kiss et al., 2013), but to our knowledge, there is no previous evidence of theta power augmentation after NMDAr blockade. The differential effect in hippocampal theta vs. PFC theta, indicates that the increase of theta power in the PFC region is not propagated, or at least not by the activity generated by the hippocampus.

The emergence of HFO in the PFC following NMDAr blockade is an intriguing phenomenon. We hypothesize that this increase is possibly produced by the alteration of the firing frequency of interneurons and pyramidal neurons and/or the desynchronization of different clusters of neurons (Homayoun and Moghaddam, 2007; Ibarz et al., 2010). On the other hand, the strong relationship between theta activity and high HFO oscillations suggests that this could be organized by the local theta.

Our results indicate that MK801 produced small changes in the dialog between the hippocampus and PFC as indicated by the frequency coherence analysis. However, paired comparisons, showed that MK801 increased the coherence in the delta band while also augmenting the coherence in the alpha band in the monopolar configuration analysis. Therefore, changes in the dialog between the hippocampus and the PFC did not affect theta oscillations, associated with spatial working memory. However, the theta/gamma modulation between the hippocampus and PFC was significantly diminished after MK801 administration, indicating a role of NMDAr in the interaction and coupling of different bands across distantly located circuit nodes. As previously mentioned, it is relevant to highlight that HFOs recorded in the PFC were not modulated by hippocampal theta after NMDAr blockade, further supporting the idea that HFOs in the PFC are locally generated and independent of hippocampal activity. The alteration of long-range modulation and the local disruption of theta/gamma activity might be caused by shared physiological mechanisms in different structures.

The circuits connecting the hippocampus and the PFC are complex. The PFC receives direct inputs from the intermediate, ventral hippocampus, the subiculum, and indirect inputs from the dorsal hippocampus through the thalamic nucleus. Therefore, the different components of the circuit might mediate different aspects of information processing. Hence, future work should aim to record from the dorsal and ventral hippocampus and subiculum, relevant thalamic nuclei as well as the PFC simultaneously (Hoover et al., 2007; Adhikari et al., 2009; Sigurdsson and Duvarci, 2016; Tamura et al., 2017).

### NMDAr blockade diminished theta/gamma co-modulation

The co-modulation of different frequency oscillations, especially theta/gamma, is key to the successful organization of neuronal ensemble activity and information processing in local and long-range networks (Sigurdsson et al., 2010; Buzsáki and Watson, 2012; Lopez-Pigozzi et al., 2016; Navas-Olive et al., 2020). MK801 administration reduced theta/gamma co-modulation in the CA1 and PFC. This decrease affected low and high gamma bands, suggesting that theta/gamma coordination depends on NMDAr normal function and most probably on PV+ interneurons (Korotkova et al., 2010; Lopez-Pigozzi et al., 2016). The observed changes in theta and gamma co-modulation activity might be associated with modifications in neuron functioning including changes in the firing rate, spike train organization, and network/neuron loss of coherence (Homayoun & Moghaddam, 2007).

### Correct and incorrect trials were specifically associated with theta/gamma comodulation levels in the hippocampus and the PFC

Previous work indicates the dialog between the hippocampus and the PFC, is necessary for spatial working memory (Sigurdsson et al., 2010; Tamura et al., 2017). We found no specific changes in the power and coherence of different oscillatory bands, linked to correct and incorrect trials, neither in the baseline or in MK801 condition. However, we observed that CA1-PFC theta/ low gamma co-modulation correlated with the performance during the baseline condition. This relationship was disrupted after MK801 administration. On the other hand, theta/high gamma co-modulation was only significantly related to performance after MK801 administration. Therefore, theta/gamma modulation might be necessary for spatial working memory, helping to coordinate local and long-range networks. These results add to previous work in which theta/gamma co-modulation range was critical for spatial working memory performance (Tamura et al., 2017).

### Different factors mediate the effects of NMDAr hypofunction

Our study found that MK801 alters the typical relationship between oscillations and motor behavior, theta/gamma co-modulation, and cognition. However, the cellular mechanisms underlying these changes need further investigation. It is likely that different cell types bear different roles in NMDAr hypofunction’s pathophysiology. This is supported by different work showing that the expression of NMDAr on pyramidal neurons is necessary for MK801’s disruption of the hippocampus and PFC interaction while NMDAr expression in interneurons appears to mediate the motor effects of MK801 (Belforte et al., 2010; Carlén et al., 2012; Bygrave et al., 2016; Hudson et al., 2020). Previous work demonstrated different interneuron types differentially affect oscillatory activity and cognitive function. Thus, PV+ interneurons partial disconnection of the excitatory-inhibitory circuits specifically affected gamma rhythms and executive function, generating an SCZ-like phenotype. On the other hand, a partial disconnection of cholecystokinin (CCK+) interneurons specifically affected theta oscillations, place cell firing, and spatial cognition (del Pino et al., 2013; del Pino et al., 2017). Similarly, we can expect a different function of NMDAr in different interneuron types. Scrutinizing these possible differences by means of opto-tagging and manipulating NMDAr activity during different conditions would help to unveil the role of NMDAr in micro and macro circuits involved in oscillatory activity, cognition, and the link between both (Royer et al., 2012; Antonoudiou et al., 2020). This manipulation should consider the dynamic expression of NMDAr subtypes during neurodevelopment, across brain regions (Monyer et al., 1994; Murillo et al., 2021) and across neuronal types (von Engelhardt al., 2015).

To conclude, we have shown that NMDAr blockade by MK801 induced positive and cognitive symptoms, altering the relationship between oscillations and locomotion as well as impairing theta/gamma co-modulation and its relationship with spatial working memory performance. We also found that drug-induced changes in oscillatory activity were brain area-specific. NMDAr’s effect on different neuronal populations, and their specific roles in different brain areas, might explain the heterogeneity of the changes we observed under MK801. Further investigating how aberrant oscillatory activity, other neuronal populations, and different NMDAr subtypes contribute to SCZ’s behavioral and cognitive deficits is necessary to understand the mechanisms underlying this disorder.

## Supporting information

Refeer to fig 5

## Author contributions

JRBM and RR designed the experiments. JRBM and PA implemented the experiments, developed and implemented the analysis toolbox. FJMP participated in the analysis of the data. JRBM, RR, and PA discussed and interpreted the results. VB and LM supported the project providing resources for its development and discussed the results. JBRM wrote the paper, and the manuscript was further reviewed and data discussed by RR and PA.

## Acknowledgments

We would like to thank Laia Serratosa Capdevila for her editing assistance. We thank Liset Menéndez de La Prida and Manuel Valero for their comments and suggestions in the initial version of the manuscript. We want to thank the support provided by the members of Victor Borrell Lab. In addition, we want to thank the technical support provided by the SEA-UMH.

## References

Housh AA, Berkowitz LE, Ybarra I, Kim EU, Lee BR, Calton JL (2014) Impairment of the anterior thalamic head direction cell network following administration of the NMDA antagonist MK-801. Brain Res Bull 109:77–87

Abekawa T, Ito K, Nakagawa S, Koyama T (2007) Prenatal exposure to an NMDA receptor antagonist, MK-801 reduces density of parvalbumin-immunoreactive GABAergic neurons in the medial prefrontal cortex and enhances phencyclidine-induced hyperlocomotion but not behavioral sensitization to methamphetamine in postpubertal rats. Psychopharmacology (Berl) 192:303–316.

Adell A, Jiménez-Sánchez L, López-Gil X, Romón T (2012) Is the acute NMDA receptor hypofunction a valid model of schizophrenia? Schizophr Bull 38:9–14

Adhikari A, Topiwala MA, Gordon JA (2009) Synchronized activity between the ventral hippocampus and the medial prefrontal cortex during anxiety.

Adler CM, Goldberg TE, Malhotra AK, Pickar D, Breier A (1998) Effects of ketamine on thought disorder, working memory, and semantic memory in healthy volunteers. Biol Psychiatry 43:811–816.

Antonoudiou P, Tan YL, Kontou G, Upton AL, Mann EO (2020) Parvalbumin and Somatostatin Interneurons Contribute to the Generation of Hippocampal Gamma Oscillations. J Neurosci 40:7668

Ban TA (2007) Fifty years chlorpromazine: A historical perspective. Neuropsychiatr Dis Treat 3:495–500

Belforte JE, Zsiros V, Sklar ER, Jiang Z, Yu G, Li Y, Quinlan EM, Nakazawa K (2010) Postnatal NMDA receptor ablation in corticolimbic interneurons confers schizophrenia-like phenotypes. Nat Neurosci 13:76–83.

Beneyto M, Meador-Woodruff JH (2008) Lamina-specific abnormalities of NMDA receptor-associated postsynaptic protein transcripts in the prefrontal cortex in schizophrenia and bipolar disorder. Neuropsychopharmacology 33:2175–2186

Bokil H, Andrews P, Kulkarni JE, Mehta S, Mitra P (2010) Chronux: A Platform for Analyzing Neural Signals. J Neurosci Methods 192:146

Bowie CR, Harvey PD (2006) Cognitive deficits and functional outcome in schizophrenia. Neuropsychiatr Dis Treat 2:531

Buzsáki G, Buhl DL, Harris KD, Csicsvari J, Czéh B, Morozov A (2003) Hippocampal network patterns of activity in the mouse. Neuroscience 116:201–211.

Buzsáki G, Watson BO (2012) Brain rhythms and neural syntax: Implications for efficient coding of cognitive content and neuropsychiatric disease. Dialogues Clin Neurosci 14:345–367

Bygrave AM, Masiulis S, Nicholson E, Berkemann M, Barkus C, Sprengel R, Harrison PJ, Kullmann DM, Bannerman DM, Kätzel D (2016) Knockout of NMDA-receptors from parvalbumin interneurons sensitizes to schizophrenia-related deficits induced by MK-801. Transl Psychiatry

Callicott JH, Bertolino A, Mattay VS, Langheim FJP, Duyn J, Coppola R, Goldberg TE, Weinberger DR (2000) Physiological dysfunction of the dorsolateral prefrontal cortex in schizophrenia revisited. Cereb Cortex 10:1078–1092.

Carlén M, Meletis K, Siegle JH, Cardin JA, Futai K, Vierling-Claassen D, Rühlmann C, Jones SR, Deisseroth K, Sheng M, Moore CI, Tsai LH (2012) A critical role for NMDA receptors in parvalbumin interneurons for gamma rhythm induction and behavior. Mol Psychiatry 17:537–548.

Chen Z, Resnik E, McFarland JM, Sakmann B, Mehta MR (2011) Speed Controls the Amplitude and Timing of the Hippocampal Gamma Rhythm. PLoS One 6:e21408

del Arco A, Segovia G, Mora F (2008) Blockade of NMDA receptors in the prefrontal cortex increases dopamine and acetylcholine release in the nucleus accumbens and motor activity. Psychopharmacology (Berl) 201:325–338.

del Pino I, Brotons-Mas JR, Marques-Smith A, Marighetto A, Frick A, Marín O, Rico B (2017) Abnormal wiring of CCK^+^basket cells disrupts spatial information coding. Nat Neurosci 20.

del Pino I, García-Frigola C, Dehorter N, Brotons-Mas JR, Alvarez-Salvado E, MartínezdeLagrán M, Ciceri G, Gabaldón MV, Moratal D, Dierssen M, Canals S, Marín O, Rico B (2013) Erbb4 Deletion from Fast-Spiking Interneurons Causes Schizophrenia-like Phenotypes. Neuron 79:1152–1168

Eyjolfsson EM, Brenner E, Kondziella D, Sonnewald U (2006) Repeated injection of MK801: An animal model of schizophrenia? Neurochem Int 48:541–546.

Fernández-Ruiz A, Oliva A, Nagy GA, Maurer AP, Berényi A, Buzsáki G (2017) Entorhinal-CA3 Dual-Input Control of Spike Timing in the Hippocampus by Theta-Gamma Coupling. Neuron 93:1213–1226.e5.

Hoover WB, Vertes RP (2007) Anatomical analysis of afferent projections to the medial prefrontal cortex in the rat. Brain Struct Funct 212:149–179.

Hakami, T., Jones, N. C., Tolmacheva, E. A., Gaudias, J., Chaumont, J., Salzberg, M., O’Brien, T. J. & Pinault, D. (2009). NMDA Receptor Hypofunction Leads to Generalized and Persistent Aberrant gamma Oscillations Independent of Hyperlocomotion and the State of Consciousness. PLOS ONE, 4 (8), https://doi.org/10.1371/journal.pone.0006755.

Hargreaves EL, Côté D, Shapiro ML (1997) A dose of MK801 previously shown to impair spatial learning in the radial maze attenuates primed burst potentiation in the dentate gyrus of freely moving rats. Behav Neurosci 111:35–48

Hertzman M, Reba RC, Kotlyarov E V. (1990) Single photon emission computed tomography in phencyclidine and related drug abuse. Am J Psychiatry 147:255–256 Available at: https://pubmed.ncbi.nlm.nih.gov/2301671/

Homayoun H, Moghaddam B (2007) NMDA receptor hypofunction produces opposite effects on prefrontal cortex interneurons and pyramidal neurons. J Neurosci 27:11496–11500

Hudson MR, Sokolenko E, O’Brien TJ, Jones NC (2020) NMDA receptors on parvalbumin-positive interneurons and pyramidal neurons both contribute to MK-801 induced gamma oscillatory disturbances: Complex relationships with behaviour. Neurobiol Dis. 134:104625. doi: 10.1016/j.nbd.2019.104625

Huettner JE, Bean BP (1988) Block of N-methyl-D-aspartate-activated current by the anticonvulsant MK-801: Selective binding to open channels Proc Natl Acad Sci U S A. 1988 Feb;85(4):1307–11. doi: 10.1073/pnas.85.4.1307.

Ibarz JM, Foffani G, Cid E, Inostroza M, Prida LM de la (2010) Emergent Dynamics of Fast Ripples in the Epileptic Hippocampus. J Neurosci 30:16249–16261

Inostroza M, Brotons-Mas JR, Laurent F, Cid E, de la Prida LM (2013) Specific impairment of “What-Where-When” episodic-like memory in experimental models of temporal lobe epilepsy. J Neurosci 33:17749–17762.

von Engelhardt J, Bocklisch C, Tönges L, Herb A, Mishina M, Monyer H (2015) GluN2D-containing NMDA receptors-mediate synaptic currents in hippocampal interneurons and pyramidal cells in juvenile mice. Front Cell Neurosci

Jackson ME, Homayoun H, Moghaddam B (2004) NMDA receptor hypofunction produces concomitant firing rate potentiation and burst activity reduction in the prefrontal cortex. Proc Natl Acad Sci USA. 2004 Jun 1;101(22):8467–72. doi: 10.1073/pnas.030845510 [Accessed June 10, 2021].

James SL et al. (2018) Global, regional, and national incidence, prevalence, and years lived with disability for 354 Diseases and Injuries for 195 countries and territories, 1990-2017: A systematic analysis for the Global Burden of Disease Study 2017. Lancet 392:1789–1858.

Jodo E (2013) The role of the hippocampo-prefrontal cortex system in phencyclidine-induced psychosis: A model for schizophrenia. J Physiol Paris 107:434–440.

Jones MW, Wilson MA (2005) Theta rhythms coordinate hippocampal-prefrontal interactions in a spatial memory task. PLoS Biol 3:1–13

Kirkpatrick B, Fenton WS, Carpenter WT, Jr., Marder SR (2006) The NIMH-MATRICS Consensus Statement on Negative Symptoms. Schizophr Bull 32:214

Kiss T, Feng J, Hoffmann WE, Shaffer CL, Hajós M (2013) Rhythmic theta and delta activity of cortical and hippocampal neuronal networks in genetically or pharmacologically induced N-methyl-d-aspartate receptor hypofunction under urethane anesthesia. Neuroscience 237:255–267

Kittelberger K, Hur EE, Sazegar S, Keshavan V, Kocsis B (2012) Comparison of the effects of acute and chronic administration of ketamine on hippocampal oscillations: Relevance for the NMDA receptor hypofunction model of schizophrenia. Brain Struct Funct 217:395–409

Korotkova T, Fuchs EC, Ponomarenko A, von Engelhardt J, Monyer H (2010) NMDA Receptor Ablation on Parvalbumin-Positive Interneurons Impairs Hippocampal Synchrony, Spatial Representations, and Working Memory. Neuron 68:557–569.

Kramis R, Vanderwolf CH, Bland BH (1975) Two types of hippocampal rhythmical slow activity in both the rabbit and the rat: Relations to behavior and effects of atropine, diethyl ether, urethane, and pentobarbital. Exp Neurol 49:58–85.

Kristiansen L V., Beneyto M, Haroutunian V, Meador-Woodruff JH (2006) Changes in NMDA receptor subunits and interacting PSD proteins in dorsolateral prefrontal and anterior cingulate cortex indicate abnormal regional expression in schizophrenia. Mol Psychiatry 11:737–747.

Kropff E, Carmichael JE, Moser EI, Moser MB (2021) Frequency of theta rhythm is controlled by acceleration, but not speed, in running rats. Neuron 109:1029–1039.e8.

Lewis DA, Curley AA, Glausier JR, Volk DW (2012) Cortical parvalbumin interneurons and cognitive dysfunction in schizophrenia. Trends Neurosci 35:57–67

Long LL, Hinman JR, Chen CM, Escabi MA, Chrobak JJ (2014) Theta dynamics in rat: Speed and acceleration across the septotemporal axis. PLoS One 9

Lopez-Pigozzi D, Laurent F, Brotons-Mas JR, Valderrama M, Valero M, Fernandez-Lamo I, Cid E, Gomez-Dominguez D, Gall B, de la Prida LM (2016) Altered oscillatory dynamics of CA1 parvalbumin basket cells during theta-gamma rhythmopathies of temporal lobe epilepsy. eNeuro 3:1–20.

Malhotra AK, Pinals DA, Adler CM, Elman I, Clifton A, Pickar D, Breier A (1997) Ketamine-induced exacerbation of psychotic symptoms and cognitive impairment in neuroleptic-free schizophrenics. Neuropsychopharmacology 17:141–150.

Meltzer HY, Stahl SM (1976) the dopamine hypothesis of schizophreniaa review. Schizophr Bull. 1976;2(1):19–76. doi: 10.1093/schbul/2.1.19.

Mena A, Ruiz-Salas JC, Puentes A, Dorado I, Ruiz-Veguilla M, De la Casa LG (2016) Reduced prepulse inhibition as a biomarker of schizophrenia. Front Behav Neurosci 10:202

Miedel CJ, Patton JM, Miedel AN, Miedel ES, Levenson JM (2017) Assessment of Spontaneous Alternation, Novel Object Recognition and Limb Clasping in Transgenic Mouse Models of Amyloid-β and Tau Neuropathology. J Vis Exp:55523

Moghaddam B, Javitt D (2012) From revolution to evolution: The glutamate hypothesis of schizophrenia and its implication for treatment. Neuropsychopharmacology 37:4–15

Monyer H, Burnashev N, Laurie DJ, Sakmann B, Seeburg PH (1994) Developmental and regional expression in the rat brain and functional properties of four NMDA receptors. Neuron 12:529–540.

Murillo A, Navarro AI, Puelles E, Zhang Y, Petros TJ, Pérez-Otaño I (2021) Temporal Dynamics and Neuronal Specificity of Grin3a Expression in the Mouse Forebrain. Cereb Cortex 31:1914–1926

Murueta-Goyena Larrañaga A., Odrioizola A.B., Gargiulo P.Á., Lafuente Sánchez J.V. (2017) Neuropathological Background of MK-801 for Inducing Murine Model of Schizophrenia. In: Gargiulo P., Mesones-Arroyo H. (eds) Psychiatry and Neuroscience Update - Vol. II. Springer, Cham. https://doi.org/10.1007/978-3-319-53126-7_25

Navas-Olive A, Valero M, Jurado-Parras T, de Salas-Quiroga A, Averkin RG, Gambino G, Cid E, de la Prida LM (2020) Multimodal determinants of phase-locked dynamics across deep-superficial hippocampal sublayers during theta oscillations. Nat Commun

Olney JW, Newcomer JW, Farber NB (1999) NMDA receptor hypofunction model of schizophrenia. J Psychiatr Res 33:523–533.

Olszewski M, Dolowa W, Matulewicz P, Kasicki S, Hunt MJ (2013) NMDA receptor antagonist-enhanced high frequency oscillations: Are they generated broadly or regionally specific? Eur Neuropsychopharmacol 23:1795–1805

Owen MJ, Sawa A, Mortensen PB (2016) Schizophrenia. Lancet 388:86–97

Richard GR, Titiz A, Tyler A, Holmes GL, Scott RC, Lenck-Santini P-P (2013.) Speed Modulation of Hippocampal Theta Frequency Correlates With Spatial Memory Performance. Hippocampus. 2013 Dec;23(12):1269–79. doi: 10.1002/hipo.22164. Epub 2013 Aug 5.

Royer S, Zemelman B V, Losonczy A, Kim J, Chance F, Magee JC, Buzsáki G (2012) Control of timing, rate and bursts of hippocampal place cells by dendritic and somatic inhibition. Nat Neurosci 15:769

Sigurdsson T, Duvarci S (2016) Hippocampal-prefrontal interactions in cognition, behavior and psychiatric disease. Front Syst Neurosci 9

Sigurdsson T, Stark KL, Karayiorgou M, Gogos JA, Gordon JA (2010) Impaired hippocampal-prefrontal synchrony in a genetic mouse model of schizophrenia. Nature 464:763–767.

Tamura M, Spellman TJ, Rosen AM, Gogos JA, Gordon JA (2017) Hippocampal-prefrontal theta-gamma coupling during performance of a spatial working memory task. Nat Commun 8.

Terrazas A, Krause M, Lipa P, Gothard KM, Barnes CA, McNaughton BL (2005) Self-Motion and the Hippocampal Spatial Metric. J Neurosci 25:8085–8096

Uhlhaas PJ, Singer W (2010) Abnormal neural oscillations and synchrony in schizophrenia. Nat Rev Neurosci 11:100–113

Tort ABL, Komorowski RW, Manns JR, Kopell NJ, Eichenbaum H (2009) Theta–gamma coupling increases during the learning of item–context associations. Proc Natl Acad Sci U S A 106:20942

Valero M, de la Prida LM (2018) The hippocampus in depth: a sublayer-specific perspective of entorhinal–hippocampal function. Curr Opin Neurobiol 52:107–114.

Vollenweider FX, Vontobel P, Øye I, Hell D, Leenders KL (2000) Effects of (S)-ketamine on striatal dopamine: A [11C]raclopride PET study of a model psychosis in humans. J Psychiatr Res 34:35–43.

Wȩdzony K, Klimek V, Goembiowska K (1993) MK-801 elevates the etracellular concentration of dopamine in the rat prefrontal cortex and increases the density of striatal dopamine D1 receptors. Brain Res 622:325–329.

Wegener N, Nagel J, Gross R, Chambon C, Greco S, Pietraszek M, Gravius A, Danysz W (2011) Evaluation of brain pharmacokinetics of (+)MK-801 in relation to behaviour. Neurosci Lett 503:68–72 Available at: https://pubmed.ncbi.nlm.nih.gov/21871531/ [Accessed June 14, 2022].

Weickert CS, Fung SJ, Catts VS, Schofield PR, Allen KM, Moore LT, Newell KA, Pellen D, Huang XF, Catts S V., Weickert TW (2013) Molecular evidence of N-methyl-D-aspartate receptor hypofunction in schizophrenia. Mol Psychiatry 18:1185–1192

Weickert TW, Goldberg TE, Gold JM, Bigelow LB, Egan MF, Weinberger DR (2000) Cognitive impairments in patients with schizophrenia displaying preserved and compromised intellect. Arch Gen Psychiatry 57:907–913.

Whishaw IQ, Vanderwolf CH (1973) Hippocampal EEG and behavior: Change in amplitude and frequency of RSA (Theta rhythm) associated with spontaneous and learned movement patterns in rats and cats. Behav Biol 8:461–484.

Young CK, Ruan M, McNaughton N (2020) Speed modulation of hippocampal theta frequency and power predicts water maze learning. bioRxiv:2020.03.31.016907 Available at: https://www.biorxiv.org/content/10.1101/2020.03.31.016907v1 [Accessed July 8, 2021].

