## Supplementary figures and images for "NMDAr Blocking by MK801 Alters Hippocampal and Prefrontal Cortex Oscillations and Impairs Spatial Working Memory in Mice"

### Refeer to fig 5

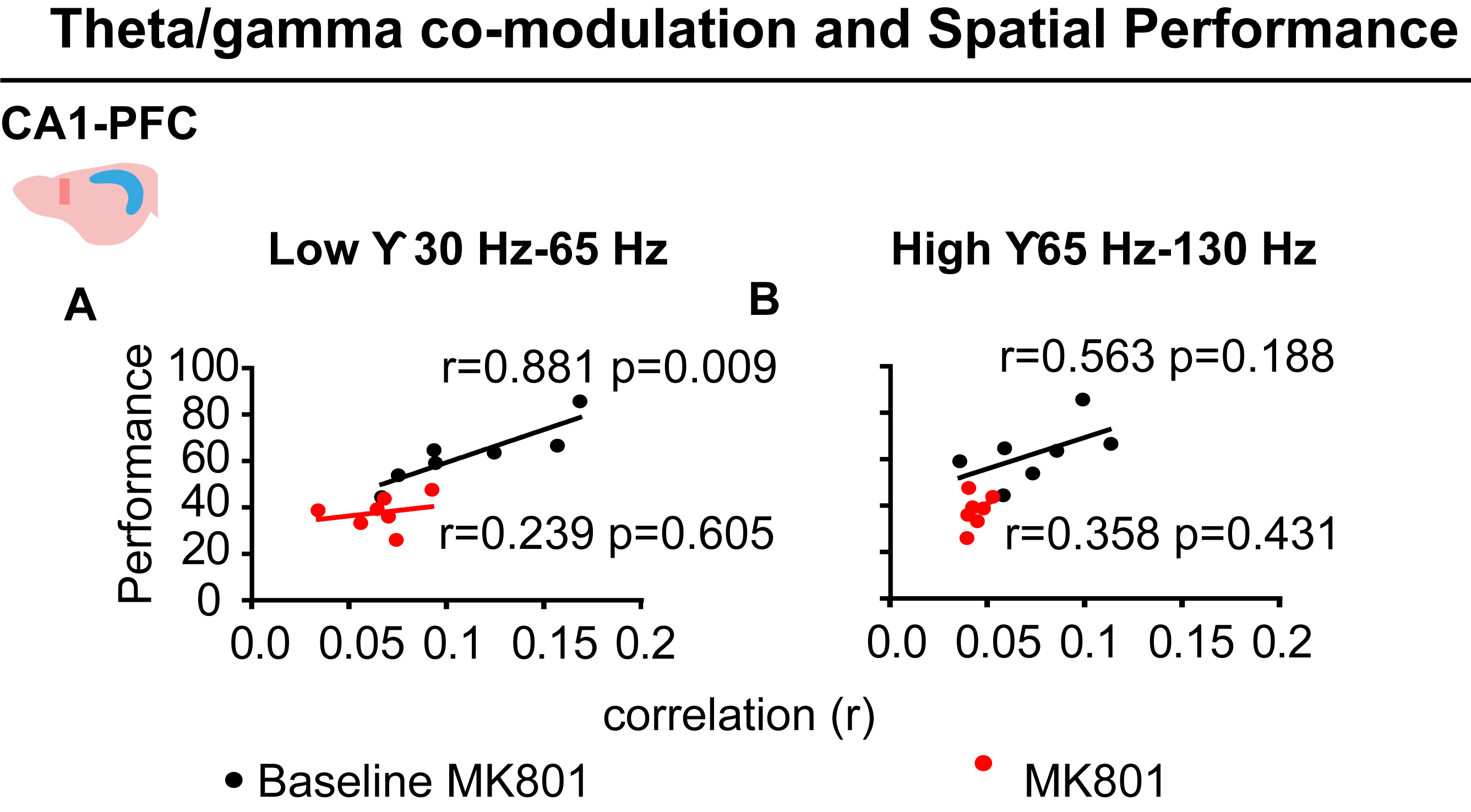
